# Characterization of primary and cultured Atlantic salmon testis cell populations using single-cell transcriptomics

**DOI:** 10.64898/2026.08.04.742726

**Authors:** Prabin Sharma Humagain, Tomasz Podgorniak, Christiaan Henkel, Victor Boyartchuk, Matthew Peter Kent, Jacob Seilo Torgersen

## Abstract

Understanding how to maintain and direct spermatogenesis *in vitro* is central to advancing reproductive biotechnologies in aquaculture species where the ability to generate gametes outside the organism could facilitate selective breeding, genetic modification, and germline preservation. However, current culture systems remain poorly defined in farmed fish species. In Atlantic salmon (*Salmo salar*), progress has been further limited by the absence of a comprehensive reference atlas of testicular cell types, making it difficult to determine how cells maintained in culture relate to their native counterparts.

To address this, we first established a single-cell RNA atlas of freshly isolated Atlantic salmon testis, resolving somatic and germ cell populations across all major stages of spermatogenesis. Primary testicular cells were then cultured under distinct conditions designed to promote either proliferation or differentiation for 14 days and subsequently subjected to single-cell RNA sequencing. To assign cell identities in cultured samples, the transcriptional profiles of cultured cells were computationally mapped onto the atlas, allowing direct comparison of cultured and native cell states.

This approach revealed condition-specific shifts in cellular composition. Media designed to promote proliferation supplemented with epidermal growth factors (EGF) and insulin-like growth factor (IGF) caused an increase in spermatogonial populations, indicating preferential support of actively dividing germ cells. In contrast, basal medium favoured the preferential survival of Sertoli cells in the absence of defined growth cues. The media expected to promote differentiation containing hormones that could stimulate male gonad development (gonadotropins and androgens) failed to do so. Further, comparative analysis of Sertoli cells across different conditions (*in vivo* and *in vitro*) revealed a loss of canonical identity markers and induction of stress-associated transcriptional programs *in vitro* compared to *in vivo*.

Together, these findings establish the first single-cell reference atlas of Atlantic salmon testis and provide a framework for evaluating and optimising testis culture systems in salmonids.

## Introduction

Spermatogenesis—the process by which motile sperm are produced from undifferentiated spermatogonial stem cells (SSC) —is a highly regulated and dynamic developmental program central to male fertility. In teleost fish, this process unfolds in a distinct architectural and physiological context compared to mammals. Rather than the continuous seminiferous epithelium characteristic of mammalian testes, fish spermatogenesis proceeds within discrete cystic units formed by Sertoli cells, a structural organisation with fundamental implications for germ cell–somatic cell communication, endocrine regulation, and aquaculture innovation (1). While the general stages of germ cell development—from spermatogonia (SPG) to spermatocytes, haploid spermatids, and ultimately mature spermatozoa—are conserved across vertebrates, fish exhibit distinct reproductive features, such as cystic spermatogenesis and species-specific endocrine regulation of germ cell development. The cellular heterogeneity within the cystic units and the molecular regulation underlying these processes are not fully understood (1,2). This includes the intermediate cell types involved in spermatogenesis in teleost fishes (3–5). Progress in this area has been hindered, in part, by the limited availability of molecular tools for validating and corelating findings from different approaches, such as transcriptomic and histological analyses, particularly in non-model organisms.

One powerful approach to investigating the molecular regulation of spermatogenesis, with potential application in aquaculture and reproductive biotechnology, is the *in vitro* culture and manipulation of SSCs. Stable spermatogonial cultures have been established in several teleost species, including zebrafish (*Danio rerio*), Honmoroko (*Gnathopogon caerulescens*), Japanese medaka (*Oryzias latipes*) and Japanese eel (*Anguilla japonica*). In these species, culture systems have supported complete spermatogenesis, producing sperm capable of fertilizing eggs and generating viable offspring (6–9). Such advances have enabled surrogate broodstock applications where donor-derived sperm are produced in host animals, germline preservation efforts, and the possibility of applying genome editing technologies in aquaculture species (10–12). However, deploying these methods to cold-water species like Atlantic salmon (*Salmo salar*) poses unique challenges, including temperature-dependent metabolic rates, differences in hormonal responsiveness, and species-specific variation in niche signalling pathways. All of which may impact the behaviour and maintenance of SSCs *in vitro* (13–15). Critically, a persistent limitation across these efforts is the absence of a well-characterised *in vivo* species-specific transcriptional reference against which to evaluate whether cultured cells genuinely recapitulate native spermatogonial states or drift toward uncharacterised transcriptional identities.

Single-cell RNA sequencing (scRNA-seq) has transformed our ability to resolve tissue transcriptional heterogeneity, enabling applications like reconstruction of developmental hierarchies or cell-type-specific gene regulatory networks at single-cell resolution. In mammals, scRNA-seq has been applied extensively to dissect the testicular microenvironment, revealing novel markers, lineage trajectories, and cell-cell interactions that govern germ cell fate (16,17). For Atlantic salmon specifically, single-cell atlases have recently been developed for only a few tissues, including skin (18), gill (19), liver (20), and spleen (21). However, to our knowledge, no such data currently exist for the testis. Given the species’ ecological and commercial importance as one of the most intensively farmed fish globally, establishing a detailed cellular reference map of the testis represents a significant and timely resource for the field.

In this study, we address this gap by first constructing the first-single-cell transcriptomic atlas of primary Atlantic salmon testicular cells, characterising the major somatic and germ cell populations and their transcriptional signatures. This atlas serves as a reference framework against which we evaluate the behaviour of early germ cells maintained under *in vitro* culture conditions designed to facilitate either proliferation or differentiation. By comparing *in vivo* and *in vitro* transcriptional profiles, we assess the extent to which each culture formulation recapitulates native spermatogonial states or supports progression through spermatogenesis, providing a principled basis for optimising culture conditions in this species.

## Methods

### Testis cell isolation

Male Atlantic salmon (n=5), each weighing approximately 200 grams were obtained from the fish facility at the Norwegian University of Life Sciences (NMBU). This size of fish was selected based on our previous observation that they consist of spermatogonial cells at different stages of development without sperm cells to be predominant. They were euthanized following Norwegian regulations for experimental animal procedures. Testes were dissected and transferred to cold Hanks’ balanced salt solution (HBSS) while blood vessels and residual blood were removed under a brightfield microscope. Testes were minced into small fragments and enzymatically digested in HBSS containing 150 U/mL collagenase at 12 °C for 2 h with gentle stirring. The resulting cell suspension was filtered through a 40 µm cell strainer and centrifuged at 400 × g for 10 min. The supernatant was discarded, and the cell pellet was washed twice with HBSS supplemented with 2% fetal bovine serum (FBS), using the same centrifugation conditions. Cells designated for single-cell RNA sequencing (hereafter referred to as the *in vivo* sample) were resuspended in HBSS and processed immediately while the cells designated for *in vitro* culture were resuspended in one of three cell culture conditions and seeded at 5X10^5^ cells per well in a 24 well tissue culture plate in 500µl of respective media. The culture media, hereafter designated as basal, proliferation and differentiation, and culture duration were formulated based on the literature on other fish species as a starting point (6,12,22,23).

### Cell culture conditions

Testis cells were cultured in triplicates for 14 days at 12°C within one of the three different media. The designation of the media is based on their expected effect rather than proven effectiveness in Atlantic salmon testis cells.

- Basal medium consisted of Leibovitz’s L-15 medium with Glutamax supplemented with 2% FBS and 1% penicillin–streptomycin.
- The other two media were prepared using Leibovitz’s L-15 medium with Glutamax as the base, supplemented with a common set of components supporting basic cell viability: 20 mM HEPES, 1% FBS, 0.5% BSA, 1X MEM non-essential amino acids, 25 µg/ml insulin (as part of ITS supplement), 200 µM adenosine, 0.1 mM beta-mercaptoethanol, 1.0 µg/ml lactic acid, 50 µM L-ascorbic acid, 1X amphotericin B, and 1X penicillin–streptomycin. This base formulation was adapted from culture conditions previously established for spermatogonial maintenance in teleost species (6,12,22,23). The two media were then distinguished by condition-specific supplements designed to promote either proliferation or differentiation.
- Proliferation medium was supplemented with epidermal growth factor (EGF, 100 ng/ml) and insulin-like growth factor (IGF, 100 ng/ml) to support spermatogonial survival and expansion (22,23), alongside progesterone at a low concentration (100 pg/ml) (12).
- Differentiation medium was designed to provide endocrine and paracrine cues supporting progression through spermatogenesis, and contained luteinizing hormone (LH, 50 ng/ml), follicle-stimulating hormone (FSH, 3.0 ng/ml), testosterone (100 ng/ml), 11-ketotestosterone (100 ng/ml), dihydroprogesterone (DHP, 50 ng/ml), progesterone (1.0 µg/ml), forskolin (50 µM), retinol (10 µM), and a mature testis extract at 50X dilution, in addition to calcium (0.03 mg/ml), magnesium (0.05 mg/ml), potassium (1.5 mg/ml), sodium pyruvate (30 µg/ml), and glutamine (1.5X) (6,12,23). Where specific concentrations had not been previously established for Atlantic salmon or closely related species, component concentrations were informed by reported physiological levels in salmonid plasma or testicular fluid or were selected based on concentrations used in comparable teleost systems. The complete composition of all three media, along with the supplier information is provided in Supplementary Table 1.

### scRNA sequencing

For single-cell RNA sequencing, the *in vivo* sample was collected in parallel with cell dissociation for culture and processed independently. Cultured cells were harvested after 14 days of culture. Culture medium containing non-adherent cells was first collected and retained. Adherent cells were then washed with PBS, and the wash fractions were pooled with the collected media to recover weakly attached cells. The remaining adherent cells were detached using TrypLE supplemented with DNase I and incubated at 15 °C for 20 min with gentle pipetting every 5 minutes. Detached cells were resuspended in basal medium and combined with the previously collected fractions. The collected cells were centrifuged at 200 × g for 5 min at 4 °C, and washed once with PBS containing 2% FBS. A final wash was performed using PBS supplemented with 1% BSA, and cells were filtered through a 40 µm cell strainer.

Cell viability was assessed using acridine orange/propidium iodide (AO/PI) staining and only samples with viability greater than 85% were used for library preparation. For each culture condition, cells from the three replicate wells were pooled prior to library preparation, resulting in one single-cell RNA sequencing library for each culture condition and one library for the *in vivo* sample. Approximately 5,000 cells per sample were loaded for library construction using the Chromium Next GEM Single Cell 3′ Kit v3.1 (10x Genomics, PN-1000269), following the manufacturer’s instructions. Libraries were prepared separately for each sample. Prepared libraries were sequenced by Novogene (Cambridge, UK) on an Illumina NovaSeq 6000 platform to a depth of 40,000 PE150 reads per cell. The raw sequencing data can be found with accession number PRJEB122847 in European Nucleotide Archive (ENA) database.

### Data analysis

Raw data were processed using the nf-core/scrnaseq (v2.4.0) (24) of the *nf-core* collection of workflows (25), utilising reproducible software environments from the Bioconda (26) and Biocontainers (27) projects. Read quality control was performed using *FastQC* (v0.11.9) (28), and gene expression quantification was carried out using *Cell Ranger* (v7.1.0) (29).

Reads were aligned to the Atlantic salmon reference genome (Salmo_salar.Ssal_v3.1.dna_sm.toplevel.withMT.fa) using a filtered gene annotation containing protein-coding genes only (Salmo_salar.Ssal_v3.1.110.filtered.withMT.gtf).

Ambient RNA contamination was estimated and removed using DecontX from the *celda* R package (v1.20.0) using default parameters (30). Cells with an estimated ambient RNA proportion greater than 50% were excluded. Then doublets were identified using both *DoubletFinder* (v2.0.4) (31) and *scDblFinder* (v1.18.0) (32), and cells classified as doublets by both methods were removed.

Downstream analysis was conducted using *Seurat* (v5.1.0) (33). Cells were filtered based on quality control thresholds, retaining cells with 200–9,000 detected genes, fewer than 40,000 transcripts, and less than 20% mitochondrial gene expression. Data were normalized using SCTransform, followed by principal component analysis (PCA). Forty principal components were used for neighbour graph construction and clustering using the Smart Local Moving (SLM) algorithm with a resolution of 0.8. Clusters were visualized using Uniform Manifold Approximation and Projection (UMAP).

Cluster-specific marker genes were identified using the Wilcoxon rank-sum test, considering genes expressed in at least 25% of cells and with a log2 fold change greater than 0.25. Cell types in the *in-vivo* dataset were assigned using combination of established marker genes and their expression patterns across clusters along with other parameters like RNA content, mitochondrial RNA percentage and number of genes in each cluster. Slingshot (v2.18.0) (34) was used with the default parameters on the subsetted spermatogenic cells only to try and infer the lineage trajectory. Cultured samples were independently integrated with the *in-vivo* reference using *Harmony* (v1.2.3) (35) with parameters theta = 10 and sigma = 0.05. Cell identities in cultured samples were inferred using a nearest-neighbour classifier in Harmony embedding space.

To assess the confidence of cell identity assignments in cultured samples, a k-nearest neighbour classifier was applied in Harmony embedding space using the *FNN* package (v1.1.4.1) (36) in R. For each query cell, the ten nearest neighbours in the reference embedding were identified, and prediction confidence was defined as the proportion of those neighbours sharing the assigned label. Cells with a prediction confidence below 0.4 were flagged as low-confidence annotations and treated with caution in downstream interpretation. Prediction confidence scores were visualised per cell using FeaturePlot and summarised across conditions using boxplots.

## Results

### Single-cell RNA sequencing and quality control

The cells from Atlantic salmon testis used for single cell RNA sequencing was a mixture of somatic cells, a high proportion of spermatogonial stem and progenitor germ cells at different stages of development, making the dataset suitable for studying spermatogenesis and germ cell maintenance *in vitro*. The dataset comprised four conditions: an *in vivo* reference sample and three *in vitro* culture conditions (designated as basal, proliferation, and differentiation media). Between ∼2,000 and ∼9,700 cells were sequenced per sample. Following quality control, including removal of ambient RNA contamination, doublets, and low-quality cells based on gene counts, transcript counts, and mitochondrial transcript proportion, a total of 16,548 cells were retained for downstream analysis (Table 1).

**Table 1.** Summary of filtering out of cells for each sample post ambient RNA estimation, doublet removal and filtering based on number of genes and transcript detected per cell, as well as mitochondrial percentage.

| S.N | Sample | Estimated number of cells | Number of cells filtered out based on ambient RNA | Number of cells filtered out as doublets | Cells filtered out based on other parameters | Number of cells after filtering |
|---|---|---|---|---|---|---|
| 1 | *In Vivo* | 9721 | - | 347 | 654 | 8720 |
| 2 | Basal | 3481 | 256 | 116 | 409 | 2700 |
| 3 | Proliferation | 2116 | 560 | 43 | 304 | 1209 |
| 4 | Differentiation | 5187 | 947 | 16 | 320 | 3904 |

Notably, cultured samples exhibited a higher proportion of cells with elevated ambient RNA contamination compared to the *in vivo* sample (Supplementary Figure 1), resulting in a larger fraction of cells being removed during filtering.

To robustly identify doublets, which arise when two cells are captured and sequenced as a single cell and can therefore confound cell-type assignment, two complementary bioinformatic pipelines were applied: *DoubletFinder* and *scDblFinder*. Each method independently classified cells as singlets or doublets, and those identified as doublets by both approaches were removed from downstream analyses, while those flagged as doublets by only one method were classified as “suspected” doublets and retained for conservative filtering.

A summary of doublet detection by each method is provided in Supplementary Table 2, while the final classification of singlets, suspected doublets, and high-confidence doublets is summarized in Supplementary Table 3. Distributions of pANN scores and doublet scores for each sample are shown in Supplementary Figure 2.

### Cell type identification in the in vivo testis sample

Unsupervised clustering using the *Seurat* package identified 24 transcriptionally distinct clusters in the *in vivo* sample of Atlantic salmon testicular cells. Cluster-specific marker genes were identified using Wilcoxon rank-sum tests (Supplementary Table 4) and used to annotate the clusters based on the expression of established marker genes, supported by additional metrics including proliferative index, mitochondrial transcript proportion, number of detected genes, and total transcript counts (Supplementary Figure 3). A summary of cluster identities, associated marker genes, and cell counts is provided in Table 2.

**Table 2.** Summary of different clusters and the markers used to define their identity in in vivo sample.

| Cluster number | Cell type | Cell number | Markers |
|---|---|---|---|
| **0** | Mid meiotic (pachytene) SPC | 1208 | *rec114, dazl, smc1b/3/4/5/6, stag3, piwil1, tdrd1, clgn* |
| **1** | Early meiotic (leptotene) SPC | 996 | *rec114, dazl, smc1b/3/4/5/6, stag3, piwil1, tdrd1, clgn* |
| **2** | Secondary SPC/RS | 986 | *adad2-rnf17, macroh2a1, topaz1, dazl* |
| **3** | Mature Sertoli cells | 892 | *amh, inha, gja1b* |
| **4** | Myofibroblast | 688 | *acta2, col1a1b, tagln,* |
| **5** | Immature sertoli cells | 410 | *amh, inha, gja1b* |
| **6** | SPG-A | 406 | *ddx4, pcna, h2az1* |
| **7** | Maturing sertoli cells | 358 | *amh, inha, gja1b* |
| **8** | Round spermatids | 327 | *adad2-rnf17, macroh2a1, topaz1, dazl* |
| **9** | SPG-B | 308 | *ddx4, pcna, ncl, h2az1* |
| **10** | Late spermatids | 265 | *nd1-5, cox1-3, cytb, atp6, atp8* |
| **11** | Early meiotic (zygotene) SPC | 258 | *rec114, dazl, smc1b/3/4/5/6, stag3, piwil1, tdrd1, clgn* |
| **12** | B cells | 247 | *ccnd2a, ebf1, bach2* |
| **13** | Immature erythrocytes | 203 | *gata1, hba4, hbba2* |
| **14** | Pericytes + Vascular endothelial cells | 179 | *rgs5a, pdgfrb, acta2, mcamb, pecam* |
| **15** | Mature erythrocytes | 165 | *hbb1, hbaa2* |
| **16** | Early meiotic (preleptotene) SPCs | 161 | *rec114, dazl, smc1b/3/4/5/6, stag3, piwil1, tdrd1, clgn* |
| **17** | Secondary spermatocytes | 146 | *dazl, adad2-rnf17, macroh2a1, topaz1, dazl* |
| **18** | T cells | 142 | *ptprc, coro1a, runx3* |
| **19** | Elongating spermatid | 117 | *adad2-rnf17, macroh2a1, topaz1, dazl* |
| **20** | Leydig cells | 93 | *star* |
| **21** | Stromal fibroblast | 64 | *col1a1, dcn, crip1* |
| **22** | Macrophages | 51 | *csf1r, cebpa* |
| **23** | Somatic-mixed | 50 | … |

Germ cell populations dominated this baseline dataset, with meiotic and post-meiotic cells collectively accounting for the majority of captured cells (Table 2). Among these, meiotic spermatocytes represented the largest proportion, spanning multiple transcriptional states corresponding to prophase I progression. In contrast, spermatogonial populations comprised a smaller fraction of the dataset, consistent with their role as a relatively rare progenitor pool (37,38). Among somatic cell types, Sertoli cells were the most abundant, represented by multiple transcriptionally distinct clusters corresponding to different maturation states, while other somatic populations, including Leydig cells, immune cells, and mesenchymal cell types, were present at comparatively lower frequencies.

To enable robust comparison of cellular composition across samples in downstream analysis, cell types were aggregated into seven broad categories based on shared biological function and lineage. These categories, used consistently across all *in vivo* and *in vitro* datasets, were: (i) erythroid cells (ii) somatic support cells (iii) immune cells (iv) mesenchymal/vascular cells (v) pre-meiotic germ cells (vi) meiotic germ cells and (vii) post-meiotic germ cells. A summary of cluster identities, associated marker genes, and cell counts is provided in Table 3, and the categories defined here were applied consistently across all samples in subsequent analyses.

i. *Erythroid cells* Two clusters (13 and 15) corresponded to erythrocytes at different maturation stages, characterised by high expression of hemoglobin genes, and were grouped into the erythroid category.
ii. *Somatic support cells* Somatic support cells category comprised primarily Sertoli cells, which formed three transcriptionally distinct but partially overlapping clusters (clusters 3, 5, and 7), reflecting a maturation continuum rather than discrete states. Immature Sertoli cells were marked by expression of *amh*, whereas mature Sertoli cells showed elevated expression of *inha* (39–41). Leydig cells formed a well-defined cluster (cluster 20) expressing *star* (42), consistent with their steroidogenic function, and were also included in the somatic support cell category.
iii. *Immune cells* The immune cell category included three clusters (clusters 12, 18, and 22) corresponding to distinct immune populations. Macrophages were identified by expression of *csf1r* (43), B cells marked by *ebf1* (44), and a T cell cluster was characterised by elevated *coro1a* expression (45).
iv. *Mesenchymal and vascular cells* The mesenchymal/vascular category comprised four cell populations (clusters 4, 14, and 21). A myofibroblast cluster was identified by expression of multiple collagen genes alongside *pdgfra*, and *acta2* (46). Stromal fibroblasts identified by canonical fibroblast markers alongside *crisp1* expression — a potentially novel association in testicular stromal cells of Atlantic salmon, given that CRISP family members are primarily characterised in the epididymis(47). A mixed pericyte/vascular endothelial cluster(cluster 14) separable into subclusters based on marker gene profiles (Supplementary Figure 5) — pericytes expressing *rgs5a*, *pdgfrb*, and *mcamb* (46), and vascular endothelial cells identified by *pecam1* expression (48).
v. *Pre-meiotic germ cells* Annotation of spermatogonial populations required the combined expression of multiple markers alongside transcriptional and metabolic features, as no single gene uniquely defined individual spermatogenic stages. Spermatogonial clusters (cluster 6 and 9) were broadly identified by expression of *ddx4*, high proliferative activity, and absence of meiotic markers. They were then further subdivided based on transcriptional and metabolic profiles. Spermatogonia A (cluster 6, Table 3; Supplementary Figure 3) displayed lower mitochondrial transcript abundance, consistent with a predominantly glycolytic metabolic state reported in early spermatogonia (49). In contrast, spermatogonia B exhibited increased mitochondrial transcript levels, higher proliferative indices, and expression of early meiotic-associated genes including *smc1*, *smc5*, and *piwil1 (*Figure 2), consistent with progression toward meiotic entry (50,51).
vi. *Meiotic germ cells* Canonical markers of meiotic stages, such as members of the SYCP family, were not robustly detected in our dataset. So, meiotic populations (clusters 0, 1, 11, and 16) were annotated based on the combinatorial expression of genes associated with synaptonemal complex formation, chromosomal cohesion, and recombination, including *rec114, smc1b, smc3, smc4, smc5, smc6, stag3, piwil1, tdrd1*, and *clgn* (2,52). Rather than forming sharply distinct clusters, meiotic cells exhibited a transcriptional continuum consistent with progressive passage through early, mid, and late prophase I. Cells corresponding to pre-meiotic and early meiotic stages displayed high proliferative activity together with expression of early meiotic genes and elevated levels of translation-related transcripts, including *eif3a, eef2b*, and ribosomal proteins. These cells also expressed *ybx1* and retained features consistent with a transitional state between differentiating spermatogonia and meiotic entry. Cells with more advanced meiotic profiles showed increased expression of genes involved in chromosomal cohesion and recombination, including *rec114, smc3*, and *stag3*, along with components of the piRNA pathway such as *tdrd1* and *piwil1*. Cells with the highest expression of *piwil1* and *clgn* were interpreted as representing later meiotic prophase-like states.
vii. *Post-meiotic germ cells* Post-meiotic cell populations (clusters 2, 8, 10, 17, and 19) were annotated using a combination of transcriptional regulators, translational control genes, mitochondrial transcript abundance, and overall transcript complexity. Secondary spermatocytes were identified by high expression of *rnf17, dazl*, *topaz1,* and *cdc20* (53–55). Transitional populations between secondary spermatocytes and round spermatids exhibited residual meiotic gene expression alongside markers associated with meiosis II, including *macroh2a2* and *adad2/rnf17* (55,56). Round spermatids were characterised in part by strong expression of EIF4EBP3L, consistent with the onset of translational repression that follows transcriptional shutdown after the secondary spermatocyte stage (57,58). Elongating and late spermatids were further distinguished by a progressive decrease in the number of detected genes and transcripts, accompanied by an increased proportion of ribosomal and mitochondrial transcripts, reflecting transcriptional silencing and metabolic specialization characteristic of late spermatogenesis (58).

The annotated clusters are visualised in a UMAP embedding of the *in vivo* sample (Figure 1A), showing clear separation of major somatic cells and germ cell lineages alongside the transcriptional continuum characteristic of meiotic progression. Meiotic stage labels and Sertoli cell cluster distinctions reflect positions along continuous developmental trajectories rather than strictly discrete cell states and should be interpreted accordingly. Figure 1B provides lineage-specific marker expression patterns, supporting cluster annotation and the interpretation of major somatic and germ cell identities, including the separation of vascular endothelial cells and pericytes. A consolidated dot plot summarizing expression of key lineage markers across major cell types is shown in Figure 2.

To assess the quality of the original cell-type annotations, the spermatogenic cell subset was reclustered independently using the selected resolution of 1. The correspondence between the original cell-type annotations and the new clusters is summarised in Supplementary Table 5. Several new clusters were strongly associated with individual original annotations, including pachytene spermatocytes, preleptotene spermatocytes, and secondary spermatocytes/round spermatids, which accounted for over 90% of the cells in their respective clusters. Other clusters contained mixtures of cells assigned to different original annotations, particularly among populations representing neighbouring spermatogenic stages. Overall, the reclustering largely preserved the broad organisation of the original annotations while revealing overlap and additional transcriptional heterogeneity among some populations. Trajectory inference using Slingshot did not resolve a single, unambiguous trajectory spanning all annotated spermatogenic stages.

Together, these results indicate that the Atlantic salmon testes analysed contain diverse germ-cell populations spanning meiotic and post-meiotic stages, supported by a somatic system comprising Sertoli and Leydig cells, immune populations, and mesenchymal and vascular cell types.

**Figure 1.**
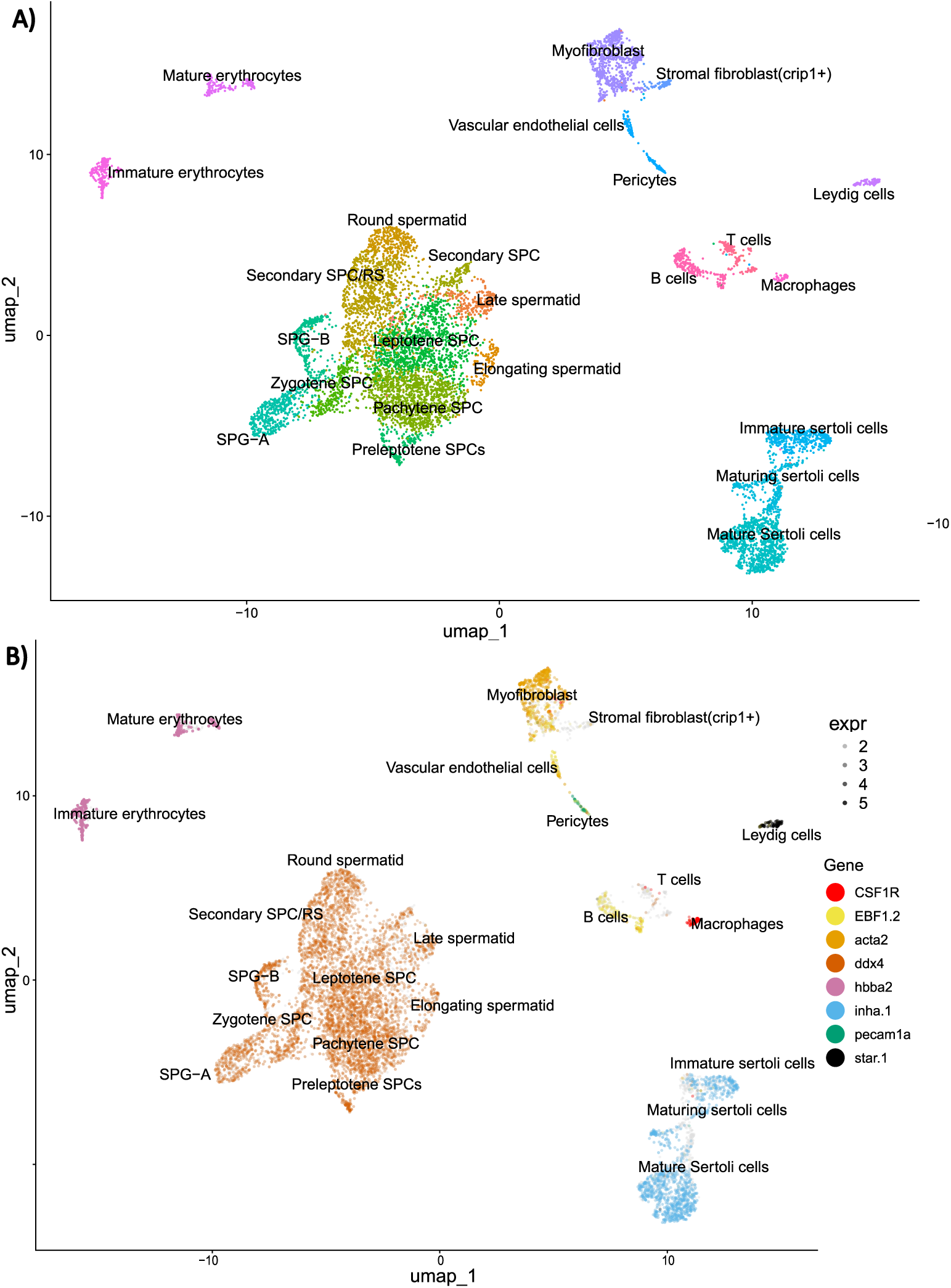
UMAP visualization of cell populations in the in vivo testis sample and expression pattern of selected marker genes. (A) Uniform Manifold Approximation and Projection (UMAP) embedding of single-cell transcriptomic data showing the distribution of germ cell and somatic cell populations. Distinct clusters corresponding to major somatic lineages (including immune, stromal, and supporting cells) are spatially separated from the germ cell clusters. Within the germ cell cluster, cells are annotated based on combinatorial expression of stage-associated genes and are organized into approximate developmental stages, including spermatogonia (SPG-A and SPG-B), preleptotene spermatocytes, and successive prophase I stages (leptotene, zygotene, and pachytene), followed by secondary spermatocytes and spermatids. These stage labels represent inferred positions along a continuous differentiation trajectory and should be interpreted as approximations rather than strictly discrete cell states. (B) Feature plots showing the expression of representative marker genes used for identification of major somatic cell types, including immune cells (e.g., CSF1R), erythrocytes (e.g., HBA/HBB), stromal and mesenchymal populations (e.g., ACTA2, PDGFRA), vascular cells (e.g., PECAM1A), and steroidogenic Leydig cells (e.g., STAR). Gene expression is overlaid on the UMAP embedding, illustrating the spatial localization and specificity of these markers across annotated clusters.

**Figure 2.**
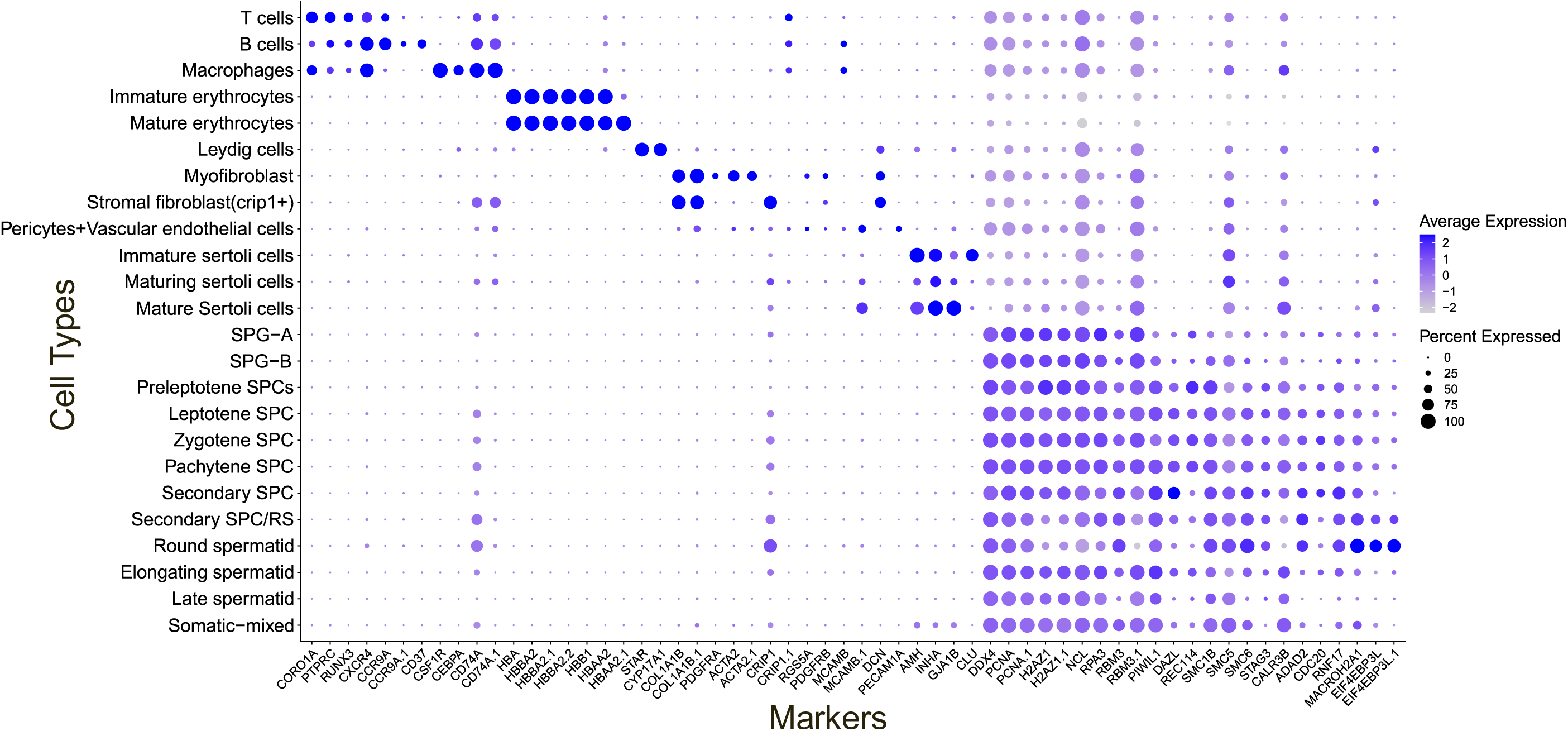
Dot plot of marker gene expression across annotated cell populations. Dot plot showing the scaled average expression (color intensity) and proportion of expressing cells (dot size) for selected marker genes across all annotated cell types. Marker genes include those associated with somatic cell populations (immune, erythroid, stromal, vascular, and Leydig cells) as well as germ cell populations spanning spermatogonial, meiotic, and post-meiotic stages. Cell-type annotation is based on the combined expression of multiple markers, and patterns of gene expression recapitulate both lineage identity and progression through spermatogenesis.

### Integration of cultured samples with the in vivo reference

To enable consistent annotation of cultured cells, each cultured dataset (basal, proliferation, and differentiation conditions) was integrated separately with the *in vivo* testis dataset using Harmony, treating the *in vivo* sample as a transcriptional reference. This approach was adopted because initial attempts to integrate all samples simultaneously revealed that culture-induced transcriptional differences dominated the joint embedding, obscuring the underlying cell-type structure and preventing reliable annotation.

For each media formulation, unsupervised clustering was first performed on the cultured sample alone, capturing distinct transcriptional states (Supplementary Figures 5, panel A, C, E). Subsequently, clustered cultured cells were integrated with the *in vivo* reference, and cell identities were inferred based on their proximity to annotated *in vivo* cell types in Harmony embedding space (Supplementary Figures 5, panel B, D, F).

To assess the robustness of cell-type annotations, prediction confidence scores were calculated for each cultured cell based on how consistently each cultured cell grouped with neighbouring reference cells of the same annotated identity following integration. Cells falling below a confidence threshold of 0.4 were considered low-confidence assignments. Across conditions, prediction confidence was generally high, supporting the reliability of reference-guided annotation (Supplementary Figure 6). However, the proliferation media showed a notably lower median confidence score and greater variance compared to basal and differentiation media, suggesting an increased transcriptional divergence from *in vivo*-defined cell states.

To assess how different culture media influenced testicular cell composition, we compared the relative abundance of major cell classes across *in vivo*, basal, proliferation, and differentiation samples (Figure 3) using the same seven biologically related categories as before.

**Figure 3.**
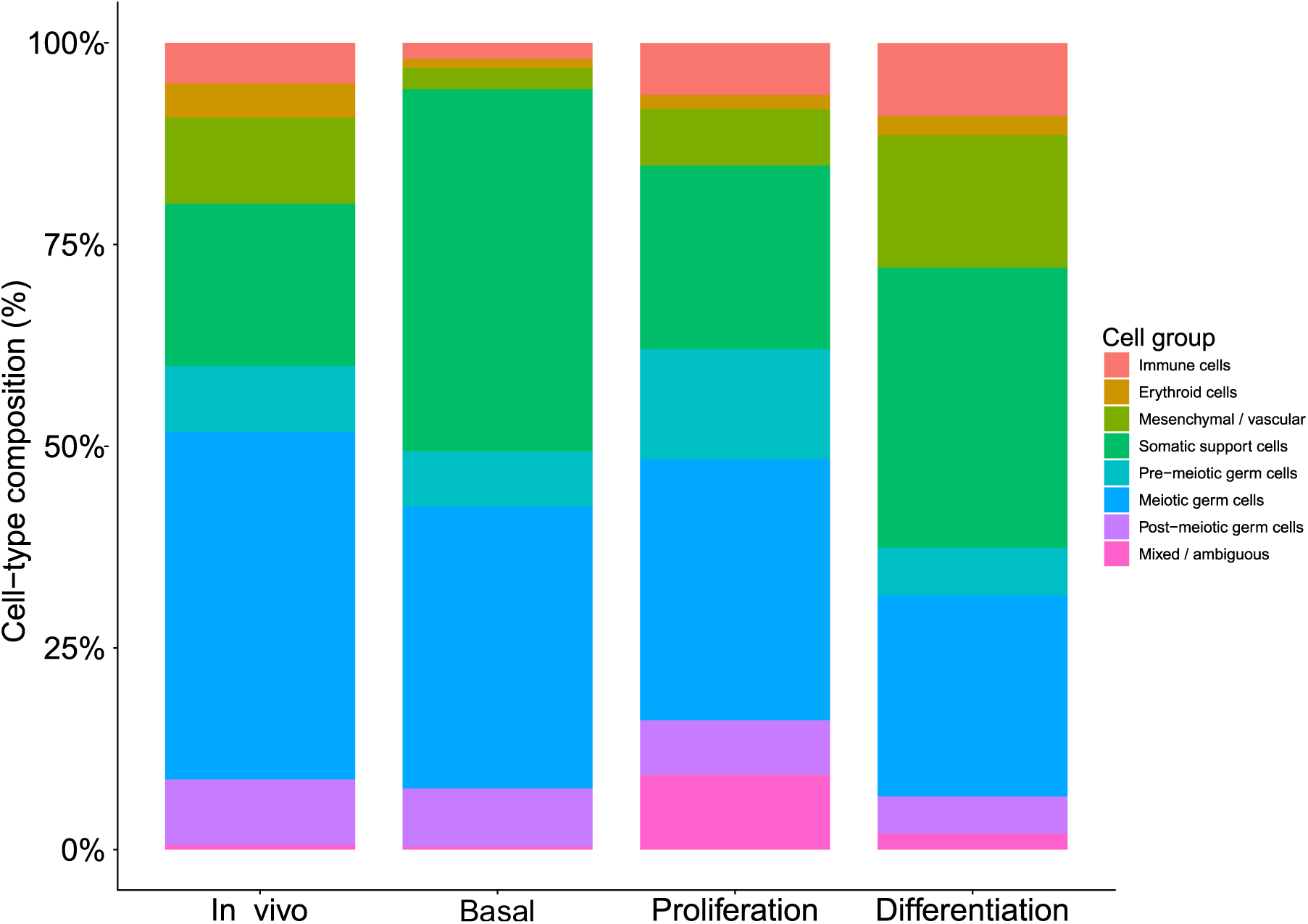
Composition of major cell groups across in vivo and in vitro conditions. Stacked bar plots showing the relative proportions of major cell groups in the in vivo testis sample and in vitro cultures under basal, proliferation, and differentiation conditions. Cell groups are aggregated into broad categories, including pre-meiotic germ cells, meiotic germ cells, post-meiotic germ cells, somatic support cells, mesenchymal/vascular cells, erythroid cells, immune cells, and mixed/ambiguous populations.

The *in vivo* sample showed a relatively even distribution of germ and somatic cell populations, with meiotic and post-meiotic germ cells representing the predominant populations, consistent with active spermatogenesis in the testis tissue. In contrast, cultured samples displayed marked shifts in cellular composition that depended on the culture media.

Cells cultured in proliferation medium showed a pronounced enrichment of spermatogonial populations, particularly SPG-A and SPG-B–derived cells, accompanied by a relative reduction in meiotic and post-meiotic germ cells. This pattern is consistent with a higher representation of cell assigned to early germ-cell populations in the proliferation sample. In contrast, basal medium was characterized by a strong enrichment of Sertoli cells, particularly mature Sertoli cells, suggesting preferential survival or expansion of somatic support cells under baseline culture conditions. Morphological observations of basal-cultured cells confirmed the presence of a heterogeneous germ cell population at 48 hours post-plating, comprising round cells of varying sizes consistent with a mixed spermatogonial population (Supplementary Figure 7).

Culturing cells in the differentiation medium did not lead to notable enrichment of late-stage germ cells. Instead, this condition resulted in increased representation of mesenchymal and somatic cell populations, with comparatively modest contributions from meiotic and post-meiotic germ cells. Together, these observations indicate that while proliferative culture conditions favour maintenance or expansion of early germ cell states, the differentiation sample did not show clear enrichment of late-stage germ-cell population under the conditions tested. A detailed breakdown of individual cell-type proportions across samples is provided in Supplementary Table 6.

To move beyond compositional shifts and examine the transcriptional consequences of culture at the gene level, differential expression analysis was performed for mature Sertoli cells, comparing *in vivo* with cultured samples. Mature Sertoli cells were selected for this analysis because they were represented in sufficient numbers across all conditions to support reliable statistical comparison, unlike most other cell types which showed pronounced compositional shifts between conditions. This analysis revealed a marked downregulation of canonical Sertoli cell markers in cultured cells — including *amh, inha,* and *gja1b*, which was reduced to near-absent levels — accompanied by large effect sizes and substantial reductions in the proportion of expressing cells (Supplementary Table 7). Concurrently, genes associated with epithelial structure and cellular stress were upregulated, including keratin family members such as *krt18b* and calcium-binding proteins such as *s100a16*. Together, these expression changes suggest a shift away from specialised Sertoli cell identity toward a more generalised stress-associated transcriptional state. Given the annotation uncertainty inherent to a non-model species, formal pathway enrichment analysis was not pursued; however, the directional consistency of these changes across individual marker genes supports the biological interpretation presented.

## Discussion

In this study, we investigated how distinct *in vitro* culture conditions influence the composition and transcriptional states of testicular cell populations in Atlantic salmon. To achieve this, we combined single-cell RNA sequencing of cultured cells with a comprehensive transcriptional reference atlas of the developing testis, enabling cell-type– resolved assessment of culture-induced changes. Through rigorous quality control, ambient RNA correction, doublet detection, and integrative analysis, we resolved a comprehensive spectrum of somatic and germ cell populations in Atlantic salmon testis. We observed differences in cell-type composition and transcriptome profiles between the freshly isolated reference and the cultured cells, with proliferative conditions favouring spermatogonial populations. We also found that our differentiation medium was insufficient to promote efficient post meiotic progression. Overall, our results provide a transcriptomic reference to enable further studies of Atlantic salmon testicular cell populations and their responses under *in vitro* culture conditions, a goal with direct relevance to reproductive management and genetic preservation in salmonid aquaculture.

### Cellular landscape of salmon testis

Our *in vivo* dataset resolved a broad spectrum of somatic and germ cell populations spanning all major stages of spermatogenesis, including spermatogonia, meiotic spermatocytes at multiple substages, secondary spermatocytes, and post-meiotic spermatids, alongside well-defined somatic populations such as Sertoli cells, Leydig cells, vascular endothelial cells, mesenchymal cells, immune cells, and erythrocytes. Cell-type annotation relied on a combination of established marker genes and complementary metrics such as proliferative index, mitochondrial transcript proportion, and transcript complexity. This integrative annotation approach was particularly important for germ cell populations, where no single marker uniquely defined individual stages, and instead reflected progressive metabolic and transcriptional remodelling during spermatogenesis. Further, this was also necessary because the majority of well characterised markers from other species were either missing or not expressed in our samples. As for the annotation themselves, even though the independent reclustering supported the broad organisation of the original annotations, some populations remained difficult to distinguish transcriptionally. Similarly, Slingshot did not resolve unambiguous trajectories. This observed overlap may reflect biological continuity between developmental stages and transcriptomic complexity during spermatogenesis, although the contribution of dataset-related limitations and analytical choices cannot be excluded. Together, these results provide an initial biologically consistent reference map of the developing salmon testis at single-cell resolution, offering a resource against which cultured or experimentally perturbed samples can be benchmarked.

### Technical challenges associated with cultured samples

Compared to the *in vivo* sample, all our cultured samples exhibited substantially higher levels of ambient RNA contamination which is consistent with increased cell stress, cell death, and RNA release during prolonged culture, particularly under conditions that do not fully support survival of all testicular cell types. Quality control thresholds were therefore selected to balance the removal of highly contaminated cells against the retention of biologically informative populations, as more aggressive filtering would have disproportionately reduced cell numbers in several cultured samples, particularly under the proliferation condition. Despite the elevated ambient RNA signal, the retained cells displayed coherent transcriptional profiles and consistent mapping to *in vivo*-defined cell identities, suggesting that meaningful biological structure was preserved following decontamination and filtering.

### Integration strategy and preservation of cell identity

Joint integration of all samples produced embeddings dominated by culture-induced transcriptional differences rather than intrinsic cell-type identities, obscuring the biological structure necessary for comparative analysis. Integrating each cultured sample separately with the *in vivo* reference resolved this issue, preserving meaningful cell-type structure while enabling direct comparison of cultured cells to their in vivo counterparts. Reference-guided integration strategies of this kind are increasingly recognised as appropriate when experimental perturbations induce strong global transcriptional shifts (59,60), and the consistent recovery of major germ and somatic cell types across conditions in the present study underscores the value of a high-quality in vivo reference as an analytical anchor.

### Culture-dependent shifts in cell-type composition

Quantitative comparison of cell-type proportions revealed clear condition-specific differences in cellular composition that primarily reflect the underlying biological properties of each medium. Proliferation media, supplemented with growth factors such as EGF and IGF, led to marked enrichment of spermatogonial populations, particularly SPG-A and SPG-B–derived cells, consistent with its intended role in promoting early germ cell survival and expansion (23,61,62). Basal media, by contrast, showed relative enrichment of Sertoli cells. This is interpretable as preferential somatic cell retention in the absence of germ cell-specific proliferative stimuli — unlike germ cells, somatic cells such as Sertoli cells do not require specialised exogenous additives to maintain basic viability and may therefore be disproportionately represented as germ cell populations decline over time. Differentiation media, which contained gonadotropins and mature salmon testis extracts intended to provide endocrine and paracrine cues supporting later spermatogenesis, did not produce a consistent or robust increase in post-meiotic germ cell populations, indicating that these conditions were insufficient to drive or sustain meiotic progression *in vitro*.

An important interpretive consideration across all conditions is that the cells are not only responding to the media components but also adapting to the *in vitro* condition. Removal of cells from the testicular environment regulated by a complex signalling to a more defined but limited media exposes cells to altered metabolic requirements and stress, evidenced by an upregulation of stress related genes in the cultured Sertoli cells compared to those in *in vivo*. The broader distribution of prediction confidence score observed in culture media, especially proliferation, also indicates a transcriptional divergence from the *in vivo* reference atlas.

The factors underlying the failure of the differentiation medium, and the broader implications of these condition-dependent compositional shifts for the design of future culture systems, are discussed in the following section.

### Biological interpretation and implications

The preferential maintenance of spermatogonial populations in proliferation media suggests that early germ cells retain a degree of plasticity and resilience *in vitro.* In contrast, progression through meiosis and spermiogenesis appears to remain tightly constrained, consistent with the view that successful spermatogenesis depends on coordinated endocrine signalling, local growth factors gradient, and direct interactions between germ cells and somatic support cells, particularly Sertoli cells (1).

The limited efficacy of the differentiation medium is likely attributable to multiple converging factors. Among these, the use of non-species-specific gonadotropins represents a probable key contributor, as gonadotropins exert their effects through cognate receptors, and differences in receptor binding affinity and downstream signalling efficiency across species may substantially reduce their biological activity in Atlantic salmon (63). In support of this, many *in vitro* spermatogenesis systems circumvent this limitation by directly applying androgens such as 11-ketotestosterone, rather than relying on gonadotropins to stimulate endogenous steroidogenesis (23). While this approach bypasses a physiologically important regulatory step, it may better approximate the steroidogenic environment required for meiotic progression when heterologous gonadotropins are unavailable. A further contributing factor is the spatial organisation of the testicular niche: the cell–cell interactions and architectural cues required for meiosis and spermiogenesis are difficult to reproduce in two-dimensional culture systems, even in the presence of exogenous hormonal signals (64,65). Importantly, these factors may not be fully independent — gonadotropins act substantially through Sertoli cells to stimulate androgen production and paracrine signalling (1), meaning that poor gonadotropin-receptor compatibility could itself contribute to the Sertoli cell transcriptional dysfunction observed here. The relative contributions of these factors cannot be resolved from the current dataset, and distinguishing between them will require systematic manipulation of individual culture components.

### Future directions

While the present findings provide important insights into the behaviour of Atlantic salmon testicular cells *in vitro*, several considerations should inform their interpretation and guide future work. Uneven cell recovery across conditions restricted the analysis of rare populations, and transcriptional silencing in haploid germ cells limits the interpretability of post-meiotic stages (66). Additionally, cell-type proportions inferred from single-cell data may be influenced by differential survival, dissociation efficiency, and capture bias (67,68). The use of consistent quality control, reference-guided integration, and conservative interpretation mitigates many of these potential sources of bias, though they cannot be entirely eliminated. Further studies incorporating biological replication and complementary validation approaches would help establish the reproducibility of the observed cell populations and clarify relationships between closely related spermatogenic stages. In particular, combining single-cell transcriptomics with morphological and protein-level evidence could help distinguish continuous developmental transitions from populations that are transcriptionally similar but biologically distinct.

Future work should prioritise improving culture systems along two complementary axes. First, the provision of species-appropriate hormonal cues, ideally salmon-derived or recombinant gonadotropins, or direct androgen supplementation as an interim strategy, should be explored to better approximate the endocrine environment required for meiotic progression. Second, and equally important, efforts to preserve Sertoli cell functionality through co-culture systems or three-dimensional architectures that better recapitulate the native testicular niche warrant serious attention, particularly given the transcriptional dysfunction observed in cultured Sertoli cells in the current study. Resolving the relative importance of hormonal versus niche-related constraints will likely require systematic experimental dissection and represents a key open question for the field. Integration with epigenomic or proteomic data may further clarify regulatory states not captured at the transcriptomic level.

## Conclusions

In summary, our study provides a first single-cell transcriptional reference of the Atlantic salmon testis and demonstrates how distinct culture conditions differentially influence testicular cell composition and transcriptional states. While early germ cells can be partially maintained *in vitro*, faithful recapitulation of later stages of spermatogenesis remains a significant challenge, likely requiring both species-appropriate hormonal signals and the preservation of somatic niche function. These findings offer important insights for the optimization of *in vitro* germ cell culture in salmonids — a goal with important implications for reproductive management in aquaculture — and establish a single-cell genomic framework that can be extended to other teleost species facing similar biological and methodological constraints.

## Funding and Acknowledgements

The authors gratefully acknowledge the internal funding support they have received from NMBU’s infrastructure financing along with financial support from AQUAGEN AS.

## Notes

### Competing Interest Statement

The authors have declared no competing interest.

### Summary of Updates

The results are the same as the original, but the interpretations have been adjusted to match the scope and clarify the limitations of the experiments performed.

