## Supplementary figures 1 - 7 for "Characterization of primary and cultured Atlantic salmon testis cell populations using single-cell transcriptomics"

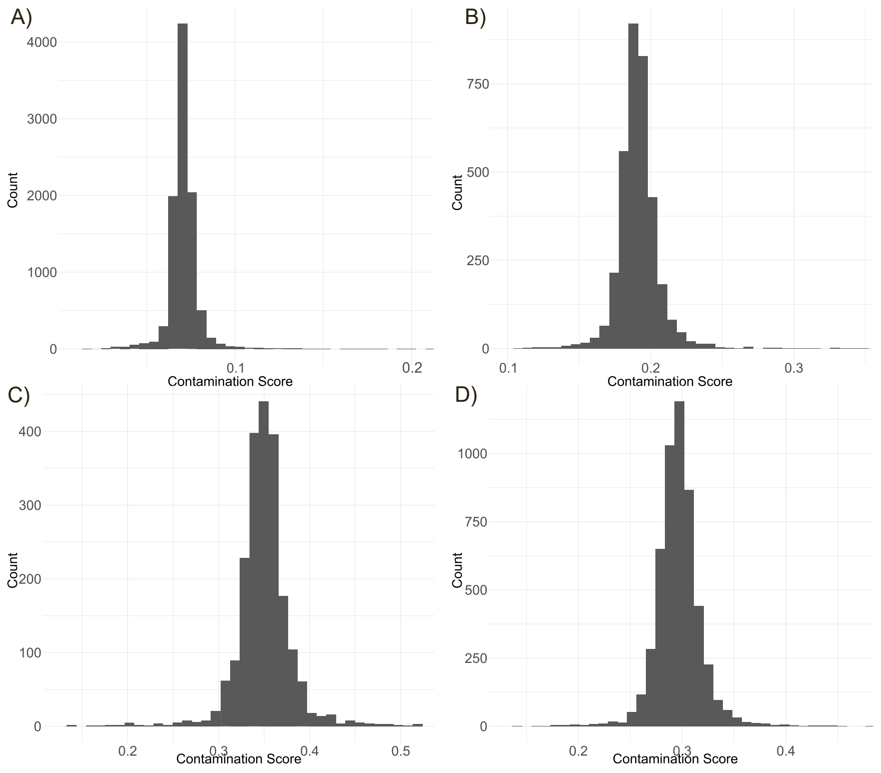


***Supplementary Figure 1*** *Contamination score of the four samples calculated using the decontX of R package celda. A) In-vivo sample B) Basal sample C) Proliferation sample D) Differentiation sample*

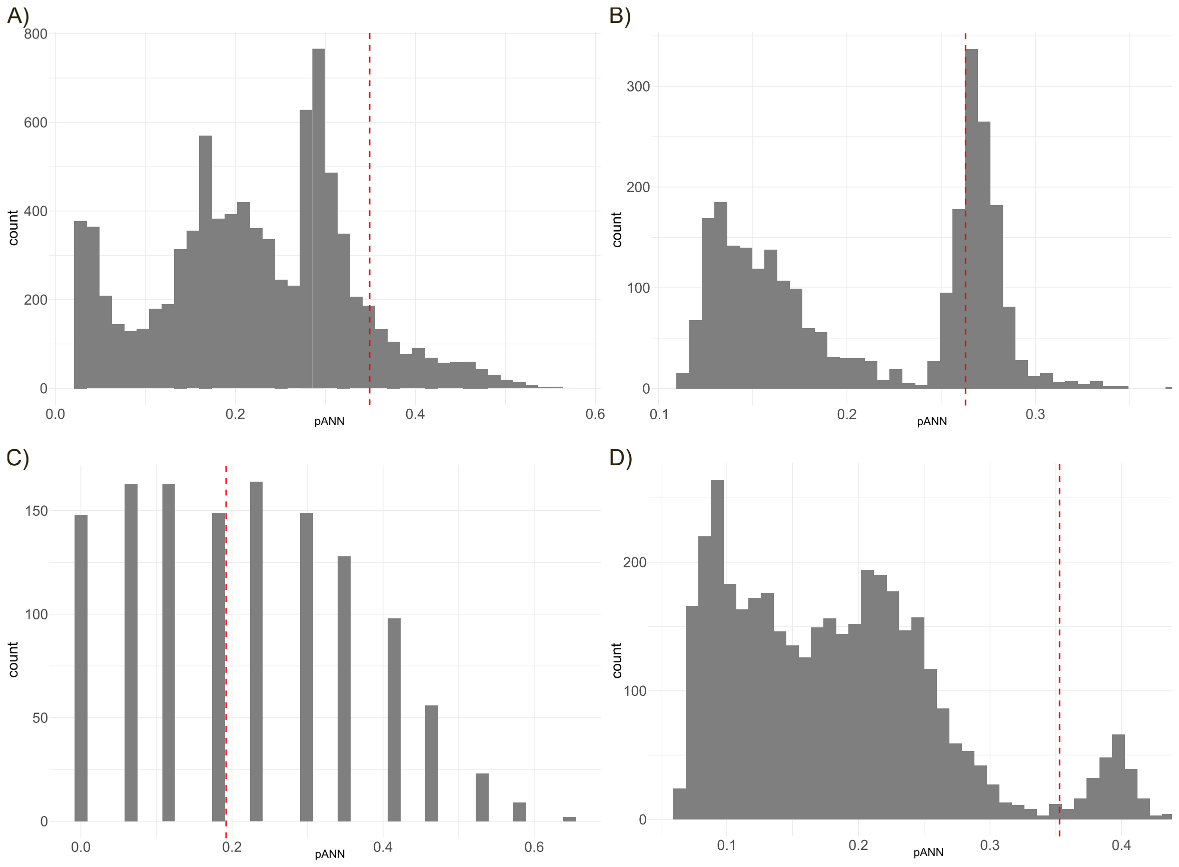


***Supplementary Figure 2*** *pANN score of different samples, calculated using the doubletfinder. The red line represents the threshold used to determine the doublets. A) In-vivo sample B) Basal sample C) Proliferation*

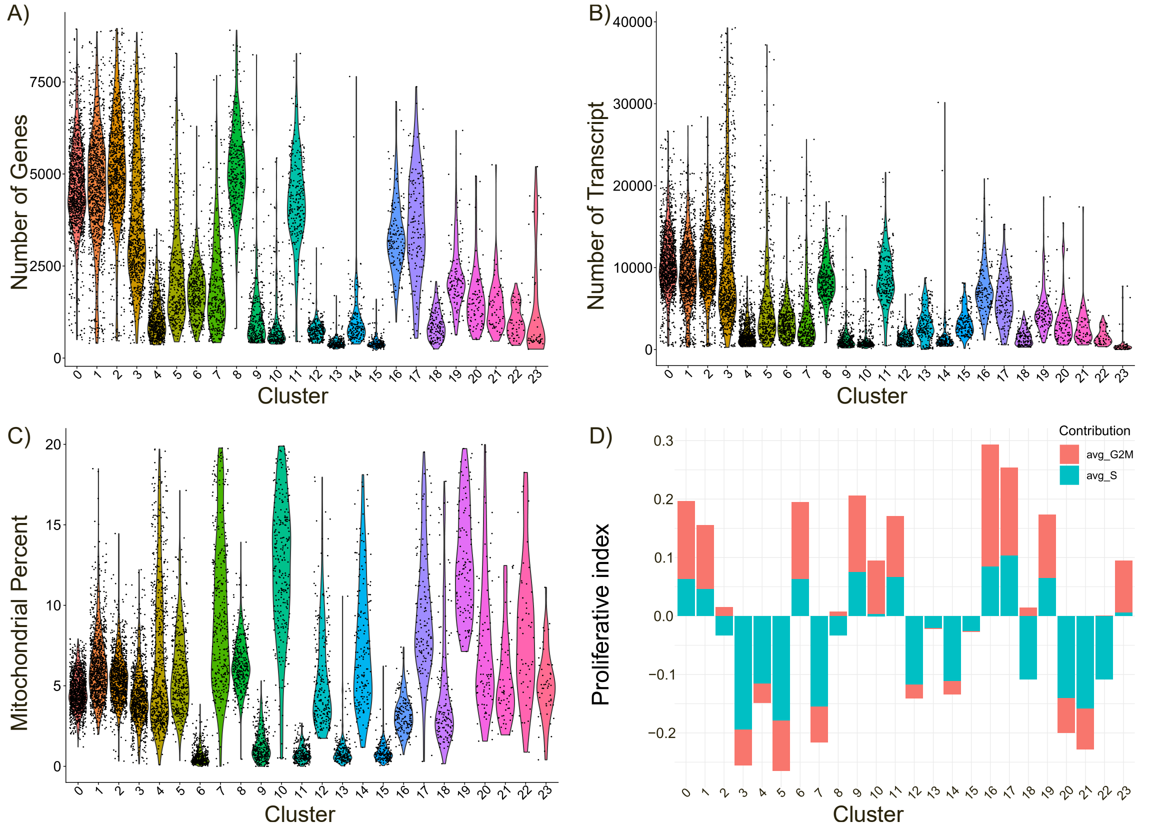


***Supplementary Figure 3*** *Metrices of different clusters of the In-vivo sample. A) Number of genes per cluster. B) Number of transcripts per cluster. C) Proportion of mitochondrial transcripts per cluster. D) Proliferative index for each cluster.*

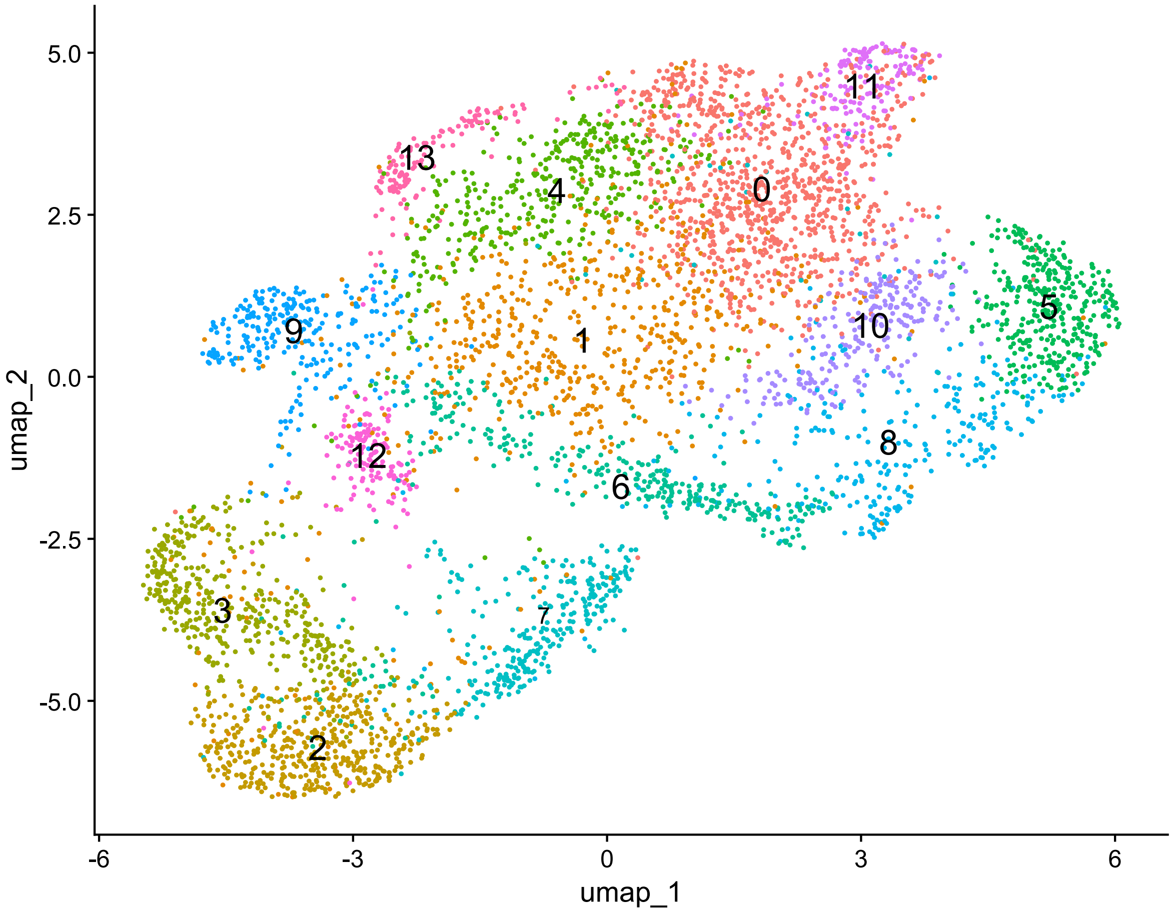


***Supplementary Figure 4 Subclustering of spermatogenic cells reveals refined structure within germ cell populations.***

*UMAP embedding of spermatogenic cells following subsetting and reclustering of germ cell populations from the full dataset. Cells are annotated based on previously assigned cell-type identities derived from the global analysis. The overall structure recapitulates the progression from spermatogonia (SPG-A and SPG-B) through meiotic spermatocytes (SPCs) to post-meiotic spermatids (round, elongating, and late stages), while providing increased resolution within the germ cell compartment.*

*Most cell populations remain consistent with the original annotations; however, some clusters, including the*

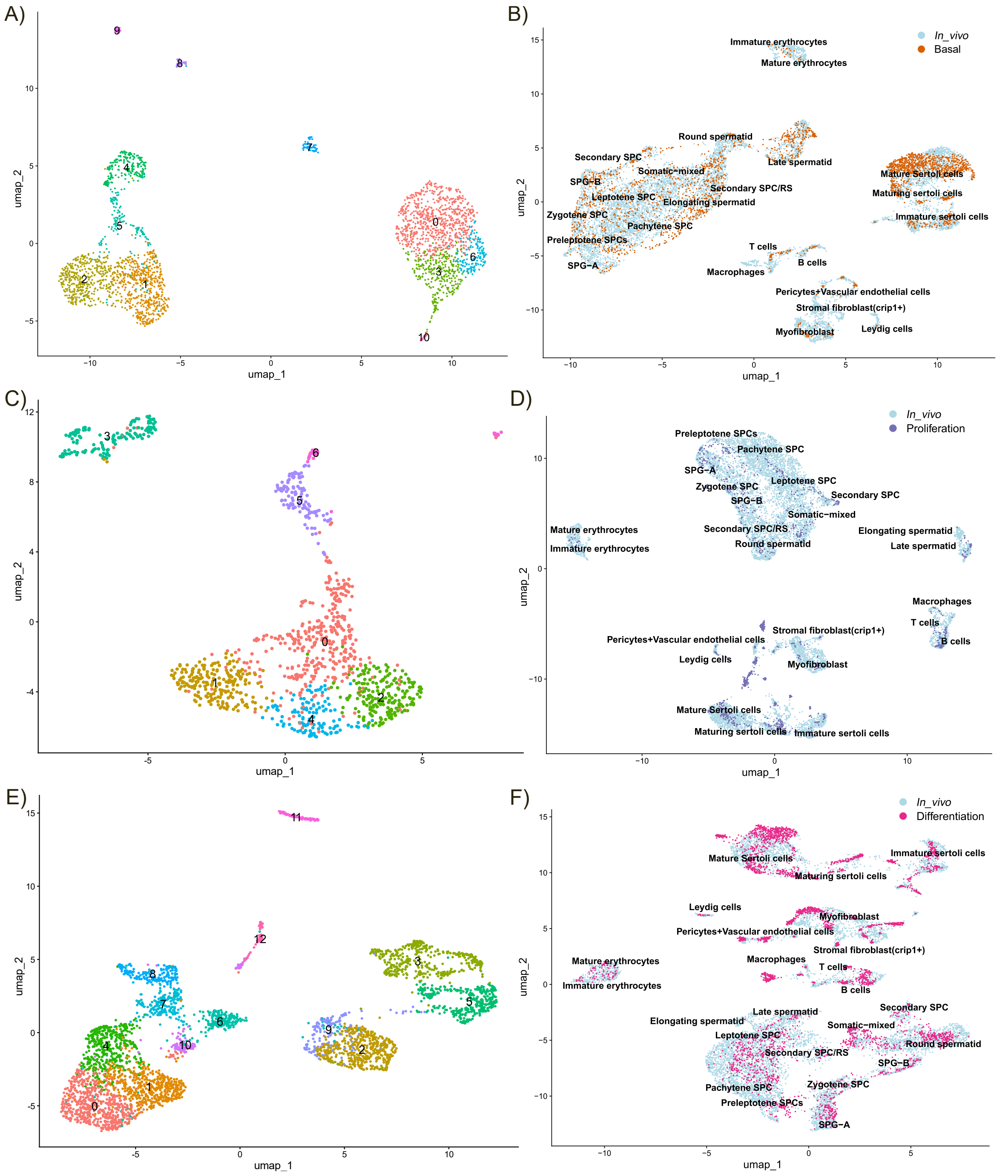


***Supplementary Figure 5 UMAP visualisation of cultured testicular cell samples before and after integration with the in vivo reference****. Panels show UMAP embeddings of cells from each culture condition prior to integration (left panels, A, C, E) and following Harmony-based integration with the in vivo reference dataset (right panels, B, D, F). In the pre-integration UMAPs, cells are coloured by unsupervised cluster identity, illustrating the transcriptional structure identified within each cultured sample in isolation. In the post-integration UMAPs, cells from the cultured sample are highlighted in colour against the in vivo reference cells, allowing direct visual assessment of how cultured cells map onto in vivo-defined cell type space. Panels A and B show the basal condition; panels C and D show the proliferation condition; panels E and F show the differentiation condition. The distinct embedding shapes observed across conditions reflect condition-specific transcriptional programs, while the integration panels illustrate the degree to which cultured cell populations correspond to native in vivo cell identities.*

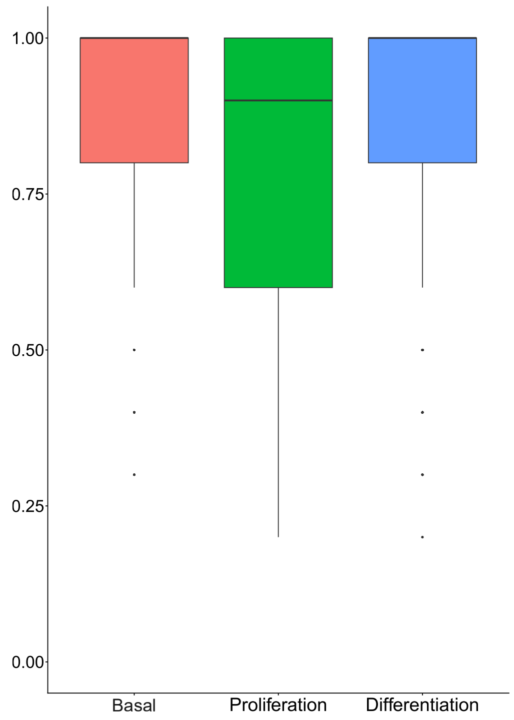


***Supplementary Figure 6*** *Distribution of prediction confidence scores for label transfer across cultured samples. Confidence values represent the proportion of nearest neighbors supporting the assigned cell type label.*

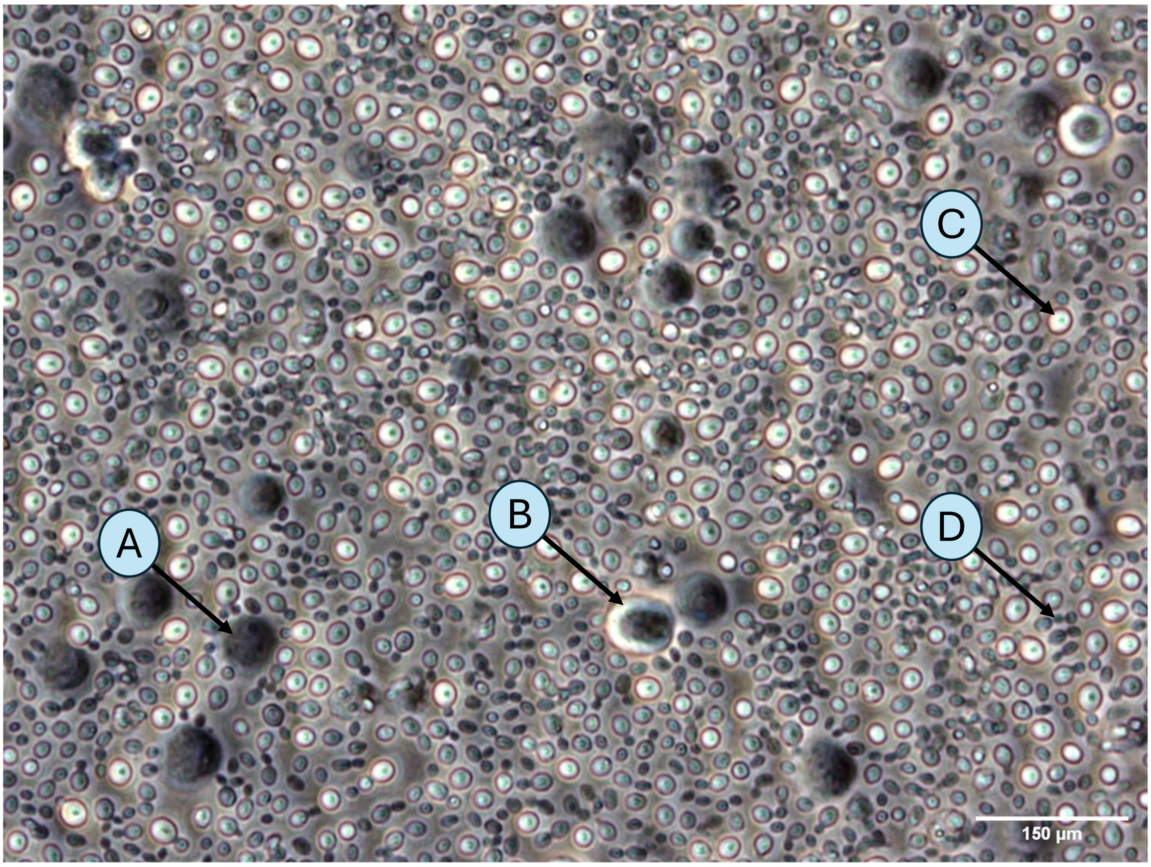


***Supplementary Figure 7. Brightfield micrograph of Atlantic salmon testicular cells in basal culture (40X objective).*** *Cells were imaged two days after plating. The image shows a heterogeneous population of round and semi-adherent cells of varying sizes, consistent with a mixed spermatogonial population. Larger, darker cells with prominent nuclei likely correspond to type A spermatogonia (A), while slightly smaller bright round cells may represent type B spermatogonia(B) or early differentiating germ cells. Much smaller bright cells observed are potentially spermatocytes(C) and darker elongating cells in chains could be spermatids(D). This image is representative of morphological observations under control conditions and is provided for qualitative context; it does not form part of the transcriptomic experiment. Scale bar not shown.*
