## Supplementary Table 1-3,5,6 for "Characterization of primary and cultured Atlantic salmon testis cell populations using single-cell transcriptomics"

***Supplementary Table 1*** *Complete composition of all culture media.*

| **Component** | **Basal medium** | **Proliferation medium** | **Differentiation medium** |
|---|---|---|---|
| Leibovitz's L-15 + Glutamax | Base | Base | Base |
| HEPES | — | 20 mM | 20 mM |
| FBS | 2% | 1% | 1% |
| BSA | — | 0.5% | 0.5% |
| MEM non-essential amino acids | — | 1X | 1X |
| Insulin (ITS) (Sigma, 11074547001) | — | 25 µg/ml | 25 µg/ml |
| Adenosine | — | 200 µM | 200 µM |
| Beta-mercaptoethanol | — | 0.1 mM | 0.1 mM |
| Lactic acid | — | 1.0 µg/ml | 1.0 µg/ml |
| L-ascorbic acid | — | 50 µM | 50 µM |
| Amphotericin B | — | 1X | 1X |
| Penicillin–streptomycin | 1% | 1X | 1X |
| EGF (Sigma, E9644) | — | 100 ng/ml | — |
| IGF (Sigma, SRP3069) | — | 100 ng/ml | — |
| Progesterone (Sigma, P8783) | — | 100 pg/ml | 1.0 µg/ml |
| Retinol (Sigma, R7632) | — | — | 10 µM |
| LH (Sigma, L6420) | — | — | 50 ng/ml |
| Testosterone (Sigma, 86500) | — | — | 100 ng/ml |
| Forskolin (Sigma, F3917) | — | — | 50 µM |
| 11-ketotestosterone (Sigma, K8250) | — | — | 100 ng/ml |
| DHP (Sigma, P6285) | — | — | 50 ng/ml |
| FSH (Sigma, F4021) | — | — | 3.0 ng/ml |
| Mature testis extract | — | — | 50X dilution |
| Calcium | — | — | 0.03 mg/ml |
| Magnesium | — | — | 0.05 mg/ml |
| Potassium | — | — | 1.5 mg/ml |
| Sodium pyruvate | — | — | 30 µg/ml |
| Glutamine | — | — | 1.5X |

***Supplementary Table 2*** *Summary of results from the two methods used to estimate doublets.*

| **S.N** | **Sample** | **DoubletFinder** | | **scDblFinder** | |
|---|---|---|---|---|---|
| | | Singlet | Doublet | Singlet | Doublet |
| 1 | *In vivo* | 7868 | 1199 | 8070 | 997 |
| 2 | Basal | 1799 | 1017 | 2645 | 171 |
| 3 | Proliferation | 623 | 629 | 1200 | 52 |
| 4 | Differentiation | 3667 | 253 | 3617 | 303 |

***Supplementary Table 3*** *Summary of the final count of doublets based on the two methods for doublet detection*

| **S.N** | **Sample** | **Combined result** | | |
|---|---|---|---|---|
| | | Singlet | Suspected Doublet | Doublet |
| 1 | *In vivo* | 8443 | 271 | 347 |
| 2 | Basal | 2503 | 197 | 116 |
| 3 | Proliferation | 1078 | 131 | 43 |
| 4 | Differentiation | 3685 | 219 | 16 |

***Supplementary Table 5*** *Correspondence between the clusters obtained by subsetting and reclustering the spermatogenic cells and the original cell-type annotations from the full dataset. Values represent the percentage of cells in each new cluster assigned to each original cell-type annotation. Percentages sum to 100% within each new cluster (column).*

| **Cell types from total data** | **New clusters after subsetting** | | | | | | | | | | | | | |
|---|---|---|---|---|---|---|---|---|---|---|---|---|---|---|
| | **0** | **1** | **2** | **3** | **4** | **5** | **6** | **7** | **8** | **9** | **10** | **11** | **12** | **13** |
| Pachytene SPC | 91.8 | 9.6 | 0 | 0 | 43.4 | 0.8 | 0 | 6.2 | 6.2 | 0 | 4.9 | 4.3 | 0 | 6.7 |
| Leptotene SPC | 6.7 | 78.1 | 0.2 | 4.3 | 53.6 | 0 | 16.8 | 5.9 | 8.3 | 19.5 | 0 | 0 | 15.3 | 6.7 |
| Secondary SPC/RS | 0.9 | 12 | 33.4 | 90.4 | 2.4 | 0 | 12.6 | 86.9 | 12.4 | 0.4 | 0.8 | 0 | 0.7 | 0.7 |
| SPG-A | 0.1 | 0 | 0 | 0 | 0 | 98.1 | 0 | 0 | 12.8 | 0 | 4.5 | 0 | 0 | 0 |
| Round spermatid | 0 | 0 | 66.4 | 4.8 | 0 | 0 | 0 | 0 | 0 | 1.1 | 0 | 0 | 0 | 0 |
| SPG-B diff | 0 | 0 | 0 | 0 | 0 | 0 | 49.1 | 0.3 | 49 | 0 | 0.4 | 0 | 0 | 0 |
| Late spermatid | 0 | 0.2 | 0 | 0.2 | 0 | 0 | 17.4 | 0 | 0 | 77.1 | 0 | 0 | 0 | 0 |
| Zygotene SPC | 0.1 | 0.2 | 0 | 0 | 0 | 0.8 | 1.2 | 0.3 | 10 | 0 | 89.4 | 0 | 0 | 0 |
| Secondary SPC | 0 | 0 | 0 | 0.2 | 0.2 | 0 | 3 | 0 | 1 | 1.9 | 0 | 0 | 84 | 0 |
| Preleptotene SPCs | 0.3 | 0 | 0 | 0 | 0 | 0.3 | 0 | 0.3 | 0.3 | 0 | 0 | 95.7 | 0 | 0 |
| Elongating spermatid | 0 | 0 | 0 | 0 | 0.2 | 0 | 0 | 0 | 0 | 0 | 0 | 0 | 0 | 85.9 |

***Supplementary Table 6*** *Summary of different types of cells in different samples*

| CellType | *In-vivo* | *In-vivo* % | Basal | Basal % | Pro. | Pro. % | Diff. | Diff. % |
|---|---|---|---|---|---|---|---|---|
| T cells | 142 | 1.63 | 35 | 1.29 | 15 | 1.24 | 184 | 4.71 |
| B cells | 247 | 2.83 | 19 | 0.7 | 57 | 4.71 | 162 | 4.14 |
| Macrophages | 51 | 0.58 | 0 | 0 | 6 | 0.5 | 7 | 0.18 |
| Immature erythrocytes | 203 | 2.33 | 13 | 0.48 | 18 | 1.49 | 31 | 0.79 |
| Mature erythrocytes | 165 | 1.89 | 18 | 0.66 | 3 | 0.25 | 62 | 1.59 |
| Myofibroblast | 688 | 7.89 | 34 | 1.26 | 39 | 3.22 | 473 | 12.1 |
| Stromal fibroblast(crip1+) | 64 | 0.73 | 0 | 0 | 11 | 0.91 | 31 | 0.79 |
| Pericytes+Vascular endothelial cells | 179 | 2.05 | 37 | 1.37 | 35 | 2.89 | 140 | 3.58 |
| Leydig cells | 93 | 1.07 | 3 | 0.11 | 12 | 0.99 | 15 | 0.38 |
| Immature sertoli cells | 410 | 4.7 | 124 | 4.58 | 38 | 3.14 | 191 | 4.89 |
| Maturing sertoli cells | 358 | 4.11 | 77 | 2.84 | 87 | 7.19 | 240 | 6.14 |
| Mature Sertoli cells | 892 | 10.23 | 1011 | 37.32 | 138 | 11.4 | 909 | 23.25 |
| SPG-A | 406 | 4.66 | 67 | 2.47 | 116 | 9.59 | 132 | 3.38 |

| | | | | | | | | |
|---|---|---|---|---|---|---|---|---|
| SPG-B diff | 308 | 3.53 | 119 | 4.39 | 50 | 4.13 | 99 | 2.53 |
| Preleptotene SPCs | 161 | 1.85 | 48 | 1.77 | 35 | 2.89 | 44 | 1.13 |
| Leptotene SPC | 996 | 11.42 | 233 | 8.6 | 66 | 5.45 | 309 | 7.9 |
| Zygotene SPC | 258 | 2.96 | 4 | 0.15 | 12 | 0.99 | 67 | 1.71 |
| Pachytene SPC | 1208 | 13.85 | 251 | 9.27 | 32 | 2.64 | 224 | 5.73 |
| Secondary SPC | 146 | 1.67 | 61 | 2.25 | 118 | 9.75 | 104 | 2.66 |
| Secondary SPC/RS | 986 | 11.31 | 350 | 12.92 | 128 | 10.58 | 228 | 5.83 |
| Round spermatid | 327 | 3.75 | 62 | 2.29 | 27 | 2.23 | 130 | 3.33 |
| Elongating spermatid | 117 | 1.34 | 13 | 0.48 | 5 | 0.41 | 1 | 0.03 |
| Late spermatid | 265 | 3.04 | 120 | 4.43 | 50 | 4.13 | 49 | 1.25 |
| Somatic-mixed | 50 | 0.57 | 10 | 0.37 | 112 | 9.26 | 77 | 1.97 |
| Total | 8720 | 100 | 2709 | 100 | 1210 | 100 | 3909 | 100 |
